# Age-related differences in working memory subprocesses decomposed by the reference-back paradigm

**DOI:** 10.1101/2024.07.04.602161

**Authors:** Zsófia Anna Gaál, Boglárka Nagy, István Czigler, Petra Csizmadia, Béla Petró, Petia Kojouharova

## Abstract

We used a data-driven approach to study the electrophysiological correlates of the working memory subprocesses revealed by the reference-back paradigm. In the absence of prior research, we focused on how aging affects the four subprocesses: *updating*, *substitution*, *gate opening*, and *gate closing*.

We conducted our experiment with 25 younger adults (M=20.17±1.47) and 23 older adults (M=67.35±4.01) using the reference-back paradigm. Significant reaction time costs were observed for all four subprocesses, but age-related differences were found only in *substitution*, which was larger in older than younger adults, indicating it as being the most vulnerable subprocess in aging.

Using difference waves, we identified event-related potential components that characterize the subprocesses we studied. Regarding *updating:* three occipital negativities between 80-180 ms, 300-400 ms, and 400-1,000 ms were observed, with only the latter range showing age group differences. Source analysis showed larger activity differences in the right frontal and temporal areas for younger adults. Regarding *substitution*: a frontal positivity between 250-600 ms emerged in younger adults, while a posterior positivity between 550-750 ms was found in older adults indicating different underlying processes supported by sLORETA results. Regarding *gate opening:* three parieto-occipital components were identified: a negativity between 150-250 ms, a positivity between 300-500 ms, and a positivity between 500-700 ms, all showing age-related differences. Regarding *gate closing*: we found an occipital negativity between 150-300 ms and a frontal positivity between 300-600 ms, neither of which changed between the age groups.

From our findings, we conclude that the process of protecting information (*gate closing*) remains stable with age, despite older adults’ sensitivity to interference. Conversely, *gate opening* is sensitive to age-related changes, likely to be resulting in different brain activity patterns during *substitution* being the updating of working memory with new information.

## 1. Introduction

Among the cognitive changes associated with aging, the most well-known and concerning is the decline in working memory [1]. Working memory plays a crucial role in maintaining and manipulating short-term information and contributing to other cognitive processes such as learning, decision-making, and problem-solving. To better understand the reasons behind the changes in working memory during aging, a possible approach is to attempt to separate its individual subprocesses and examine them independently, along with the underlying brain structures. This allows us to investigate whether one or more subprocesses are affected during aging and which ones play a role in the deteriorated function of working memory.

A potential method for this exploration is provided by the reference-back paradigm developed by Rac-Lubashevsky and Kessler [2]. In this working memory model the central element is a gating mechanism which regulates the protection and updating of content in the working memory. The reference-back paradigm allows for the separation of the subprocesses of working memory – *updating*, *substitution*, *gate opening*, and *gate closing* – by manipulating the state of the gate and the need to update in the individual trials. This is a modified version of the n-back paradigm: letters (X and O) are presented randomly in succession within either a red or a blue frame, and the task is to determine whether the currently presented letter is identical or not to the last letter presented within a red frame. So, trials with a blue frame (comparison trials) require only a matching decision (the gate is closed); whilst trials with a red frame (reference trials) also require the updating of the content in working memory (the gate is open). Behavioural results indicate the following: 1) an *updating cost* with reaction times being slower for reference compared to comparison trials; 2) a *substitution cost* where reaction times are much slower for “different” compared to “same” responses in reference trials to comparison ones; 3) a *gate opening cost* where reaction times are slower for reference trials following a comparison trial (switch) than reference trials following another reference trial (no-switch); and 4) a *gate closing cost* where reaction times are slower for comparison trials following a reference trial (switch) than comparison trials following another comparison trial (no-switch). These results confirmed the separability and examinability of the presumed subprocesses [2,3]. The above subprocesses, according to the authors, can be matched with the biological processes hypothesized by the prefrontal cortex basal ganglia working memory model (PBWM [4–6]). The basal ganglia serve the gating function, separating sensory input in the sensory cortex; or internally stored input in long-term memory from the working memory representation maintained in the dorsolateral prefrontal cortex. In the baseline state, the gate is closed, preserving the content of working memory. However, when required by the task, the gate opens, allowing for the working memory to update with new information. By this the inhibitory input from the basal ganglia to the thalamus, which has bidirectional excitatory connections with the frontal cortex, is released. This mechanism enables control over working memory content: flexibly alternating between either protecting or refreshing it with new information.

The assumed biological basis of the model, the involvement of the structures mentioned above in working memory processes, is supported by fMRI results. Nir-Cohen and her colleagues [3] employed a reference-back task using faces instead of letters. Their results corroborated the involvement of the basal ganglia-thalamus-prefrontal cortex circuits with some refinements. The basal ganglia (caudate, putamen and pallidum) exhibited increased activity for *gate opening* and *substitution*, but not for *gate closing* and *updating*. Enhanced thalamic activity was identified solely for *gate opening*. The entire frontoparietal network, including the dorsolateral prefrontal cortex, the medial prefrontal cortex, and the posterior parietal cortex, was activated during *gate opening*; while a lateralized activity in the left parietal and frontal cortex was observed during *substitution*. *Updating* was associated with activity increase in the posterior parietal cortex. Pronounced activity was also noted in the fusiform face area during *gate opening* and *substitution*. In summary, the four subprocesses were linked to distinct neural activity patterns. While *gate opening* showed an activation pattern supporting the PBWM model, these structures were not involved in *gate closing*. Additionally, other structures, such as the posterior parietal cortex and fusiform face area, were also involved in working memory *updating*. However, a reanalysis of these results [7] revealed frontal, striatal, and thalamic activity during *gate closing* when the current stimulus differed from the one held in working memory, and no activity increase was observed except when the current stimulus and the one held in working memory were identical. Thus, active gate closure occurred only when it was necessary to shield the information, and not by default. Also, selective gating was observed for *gate opening*, occurring only when genuinely necessary and not during task repetition. Another study has suggested that similar neural patterns are activated regardless of whether working memory is updated with declarative or procedural content [7].

While fMRI allows us to explore which brain regions are involved in individual subprocesses, the event-related potential (ERP) method enables us to examine their temporal dynamics with millisecond precision. To date, only two studies [8,9] have investigated the reference-back task in this manner. Rac-Lubashevsky and Kessler [8] focused solely on the P3b component. The reasoning for this is that the P3b component is associated with updating the content of working memory [10]; its amplitude increases when a stimulus is relevant to the current task, yet unexpected. However, Rac-Lubashevsky and Kessler [8] concluded that the P3b was not associated with working memory updating; but rather indicated target categorization. Although Csizmadia and her colleagues [9] identified several components that played a role, they focused only on gate opening, and specifically on the differences between groups with different levels of divergent and convergent thinking.

Nevertheless, it is of note the authors in these studies drew their conclusions from the original ERPs, even though their primary concept for distinguishing working memory subprocesses relied on subtracting specific trial types. When working with ERPs, utilizing difference waves is proven to be more effective in isolating processes that exhibit distinct activity between two (or more) trial types. This approach helps to eliminate patterns that show no variation between these trial types, while emphasizing the relevant components that result from their distinct activation patterns. Consequently, this method enables us to make specific conclusions based on the understanding of how these subprocesses are uniquely activated in different trial types [11]. Therefore, in the current study, we employed difference waves, and we used a data-driven approach (cluster-based permutation t-tests) to identify the components unique to each subprocess. In this regard, our study was exploratory in nature.

In addition to exploring which subprocesses manifest in each component, we aimed to understand how these subprocesses differ between young and older adults. Taking into consideration that the macrostructure, microstructure, and neural connectivity of each involved structure change with aging (basal ganglia [12]; thalamus [13,14]; prefrontal cortex [15,16]); it is expected that individual subprocesses will also exhibit differences between young and older adults, primarily in *gate opening* and *substitution*. This is because these subprocesses show the most significant activity increase in these structures. We also applied source localisation analysis to identify possible differences in the sources of the subprocesses between younger and older adults. Any differences in the ERPs can reflect differences in the use of brain resources, e.g. the employment of compensatory mechanisms [e.g., 17]; and source localisation can help to identify the brain structures responsible for these differences.

To summarize, our study investigated the subprocesses involved in working memory and also their temporal course in the reference-back paradigm. Our aim was to identify the components of each subprocess using a data-driven approach, as earlier studies have neither done this; nor have utilized difference waves measurement. Furthermore, we endeavoured to examine how these subprocesses may change with aging.

## 2. Material and methods

### 2.1. Participants

Two age-groups participated in the experiment. There were 29 participants in each group. Five participants in the older group did not complete the session, and a further one was excluded because of a lack of valid epochs in one of the conditions. Two participants in the younger group were excluded because of recording issues, and three others were excluded because of too few epochs in some of the conditions. In the final sample, the younger group consisted of 24 participants (mean age: 20.17, SD = 1.47, range 18–23 years; 16 women; 2 left-handed), and the older group consisted of 23 participants (mean age: 67.35, SD = 4.01, range: 60–74 years; 14 women; 1 ambidextrous). To exclude dementia-related differences between the age groups, we measured the four major components of intelligence with four subtests of the Hungarian version of Wechsler Adult Intelligence Scale (WAIS-IV [18]) representing the four major components: Similarities – verbal comprehension; Digit Span – working memory; Matrix Reasoning – perceptual reasoning; and Coding – processing speed. The scaled scores (where the age-group average is 10) achieved by the younger group were as follows: Similarities: M = 11 (SD = 2.3); Digit span: M = 8.7 (SD = 2.4); Matrix Reasoning: M = 10.3 (SD = 1.8); and Coding: M = 11.3 (SD = 2.8). The older group achieved the following scores on the four subtests: Similarities: M = 13.5 (SD = 2.0); Digit Span: M = 11.4 (SD = 3.4); matrix reasoning: M = 12.8 (SD = 2.9); and Coding: M = 14.2 (SD = 3.0). Every participant had normal or corrected-to-normal vision and had no history of any kind of neurological or psychiatric disorder. All participants were paid for their contribution and were recruited between October 2018 and March 2019. The protocol was approved by the Joint Psychological Research Ethics Committee (EPKEB, Hungary) and a written informed consent was obtained from all participants.

### 2.2. Stimuli and procedure

Each participant was seated in a comfortable chair in a sound-attenuated and electrically shielded chamber. Stimuli presentation and behavioural data collection were conducted with MATLAB R2014a [19]. The stimuli were presented on a 21.5-inch LCD monitor (Asus VS229na, 60 Hz refresh rate) at a 1.4 m distance. The present study was composed of one session that included a forced-choice task; a reference-back task; a reference-back task with distractors; and a memory task where participants had to indicate if they remembered the distractor stimuli. The present paper considers only the reference-back task, and its experimental design is shown in Figure 1. The stimuli (letter “X” at 0.35° visual angle; or “O” at 0.4° visual angle) were presented in 4 pixel-wide lines against a grey background (RGB (0.45,0.45,0.45)) within either a red or a blue frame at 1.15° visual angle. Each trial started with a fixation-cross that was presented for 1,400 ms (with +/-100 ms jitter in 16.6 ms steps), followed by one of the stimuli for 500 ms (with +/-50 ms jitter in 16.6 ms steps). The trials were presented in a randomized order in blocks. Each block contained 80 trials, and nine blocks were presented for a total of 720 trials. The experimental blocks were preceded by 4 practice blocks (15 trials per block). Throughout the experimental blocks, participants were asked to keep their gaze on the centre of the screen. The task had two trial types: in reference trials (probability 0.5) the letters appeared within a red frame, and in comparison trials (probability 0.5) they were presented within a blue frame (see: Figure 1). A reference trial could follow a comparison trial (switch reference, the gate opens, switch trial) or another reference trial (no-switch reference, the gate remains open, no-switch trial). Similarly, a comparison trial could follow a reference trial (switch comparison, the gate closes, switch trial) or another comparison trial (no-switch comparison, the gate remains closed, no-switch trial). Within a block the probability of a no-switch trial was 0.75, and the probability of a switch trial was 0.25. In each trial, participants had to select whether the stimulus was or was not the same as the one in the last seen reference trial (red frame). “Same” responses, on a modified computer keyboard with only two keys, were indicated by pressing the ‘L’ key with the right index finger; and “different” responses, by pressing the ‘A’ key with the left index finger. Within all blocks of trials an equal number of “same” and “different” responses were required. The participants had a time-window 2,000 ms in which to respond. After each block, participants received feedback on their performance, including: the average reaction time; the number of correct responses; and the number of errors. Responses below 150 ms were considered invalid, and excluded from data analysis (a total of 62 trials for the experiment).

**Figure 1.**
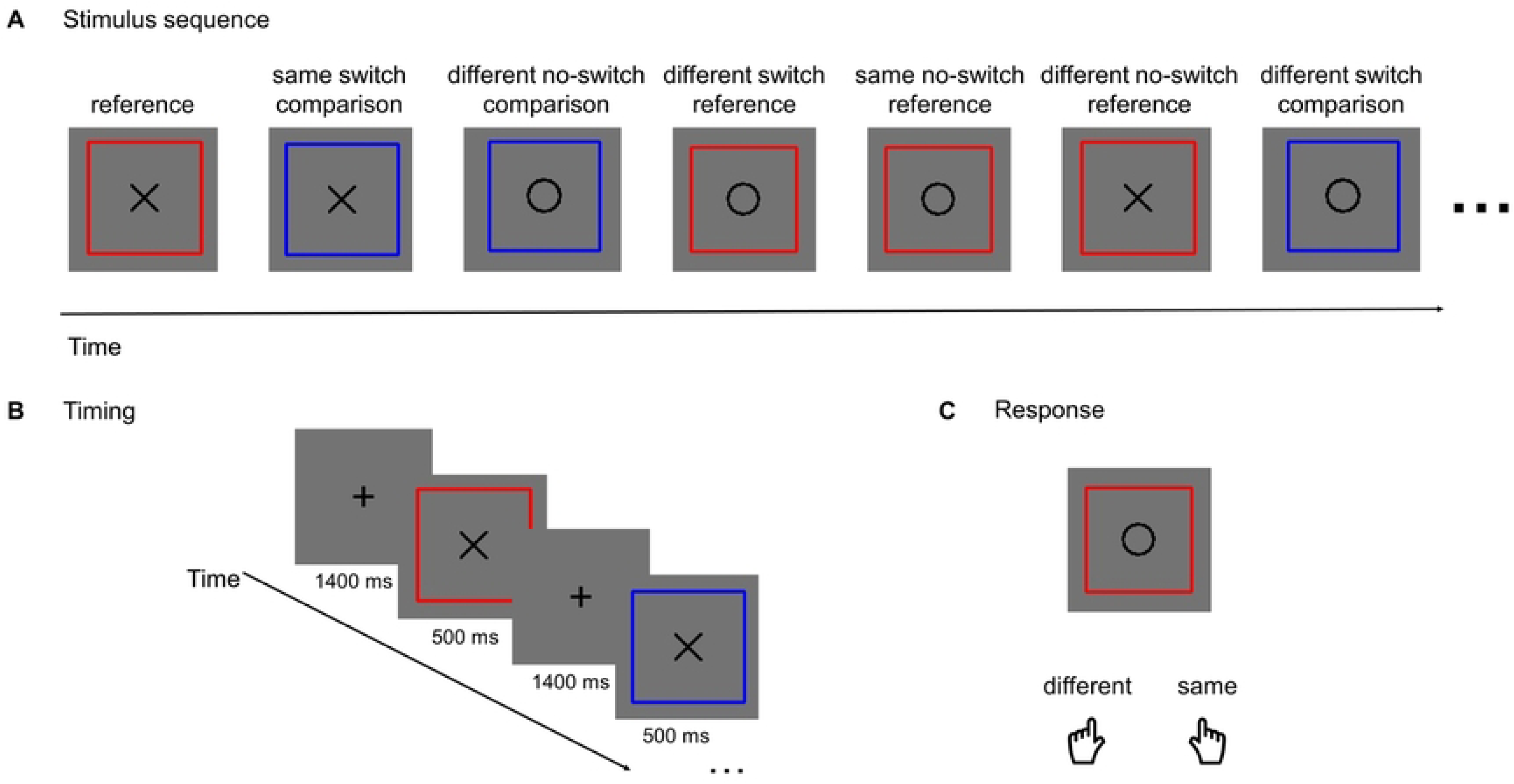
**A** A stimulus sequence example consisting of reference trials (red frame) and comparison trials (blue frame). The target stimulus was the letter X or the letter O displayed in the centre of the frame. **B** The stimulus duration was 500 ms (±50 ms). The inter-stimulus interval (ISI) duration was 1400 ms (±100 ms) during which a fixation cross was visible in the centre of the screen. **C** A response was given by pressing the left key if the target letter in the trial was different from the reference, and the right key if the target letter was the same.

### 2.3. ERP recording

EEG was recorded with BrainVision actiCHamp amplifier (Brain Products GMBH, DC- 280 Hz, sampling rate: 1,000 Hz). Brain activity was recorded from 32 Ag/AgCl active electrodes, in accordance with the extended 10–20 system (F7, F3, Fz, F4, F8, FC3, FC4, T7, C3, Cz, C4, T8, CP5, CP6, P7, P3, Pz, P4, P8, PO7, PO3, POz, PO4, PO8, O1, Oz, O2, with AFz as the ground) using an elastic electrode cap (EasyCap, Brain Products GMBH). The reference electrode was placed on the tip of the nose. Horizontal EOG was recorded with a bipolar configuration between electrodes positioned lateral to the outer canthi of the eyes (one electrode on each side). Vertical eye movement was monitored with a bipolar montage between two electrodes, one placed above and one below the left eye. The impedance of the electrodes was kept below 10 kΩ.

### 2.4. Data analysis

#### 2.4.1. Computation of the subprocesses

The subprocesses were computed as follows: 1) *updating* = no-switch reference – minus – no-switch comparison, where information is entered into working memory independent of whether it is the same as the maintained information or not; 2) *substitution* = (different no-switch reference – minus – same no-switch reference) – minus – (different no-switch comparison – minus – same no-switch comparison), where the information in working memory needs to be replaced only if the reference changes; 3) *gate opening* = switch reference – minus – no-switch reference, where the gate to working memory needs to be opened opposite to being maintained open; 4) *gate closing* = switch comparison – minus – no-switch comparison, where the gate needs to be closed opposite to being maintained closed. These computations are illustrated in Table 1. In our analysis, these computations are applied for comparing the size of the behavioural effects between the age groups as well as for calculating the difference ERP potentials.

**Table 1.** Calculation of the subprocesses in the reference-back task.

| TRIAL TYPE | Reference |  |  |  | Comparison |  |  |  |
| --- | --- | --- | --- | --- | --- | --- | --- | --- |
| GATE SWITCH | No-switch |  | Switch |  | No-switch |  | Switch |  |
| RESPONSE | Same | Different | Same | Different | Same | Different | Same | Different |
| Gate opening | - | - | + | + |  |  |  |  |
| Gate closing |  |  |  |  | - | - | + | + |
| Substitution | - | + |  |  | + | - |  |  |
| Updating | + | + |  |  | - | - |  |  |

#### 2.4.2. Behavioural data analysis

First, we calculated overall mean performance (percentage of correct responses of all trials) and overall mean reaction time (for correct responses only) to assess whether there was an overall difference between the age groups in the task. Because of the 2,000 ms time limit, outlier RTs were not removed (under 1% of RTs were between 1,500 ms and 2,000 ms in either age group). While slower reaction time (RT) was expected in the older group, we wanted to ensure that the level of performance was similar in the groups. Furthermore, in a task with a time limit there are two possible types of error: either an incorrect response is given within the allowed time limit, or there is no response (miss). So, overall incorrect response rate and misses rate were also analysed.

Mixed analyses of variance (ANOVAs) on the mean RT results for the correct responses were performed to explore the presence of the investigated effects as well as possible differences between the age groups. Only RTs for correct responses were included in the averages (see Table 1 for the factors and levels included in each ANOVA). For the *updating* subprocess, we conducted an ANOVA with TRIAL TYPE (no-switch reference, no-switch comparison) as the within-subject factor and AGE GROUP (younger, older) as the between-subject factor. A statistically significant main effect of TRIAL TYPE means presence of the *updating cost*, while a significant TRIAL TYPE × AGE GROUP interaction means a difference in the *updating cost* between the age groups. For the *substitution* subprocess, we conducted an ANOVA with TRIAL TYPE (no-switch reference, no-switch comparison) and RESPONSE (same, different) as the within-subject factors and AGE GROUP (younger, older) as the between-subject factor. A significant TRIAL TYPE × RESPONSE interaction shows the presence of *substitution*, while a significant TRIAL TYPE × RESPONSE × AGE GROUP interaction shows a difference in the subprocess between the age groups. For the *gate opening* subprocess, we conducted an ANOVA with GATE SWITCH (switch reference, no-switch reference) as the within-subject factor and AGE GROUP (younger, older) as the between-subject factor. A significant main effect of GATE SWITCH indicates the presence of *gate opening*, while a significant GATE SWITCH × AGE GROUP interaction supports an assumption for an age-based difference in the subprocess. Lastly, for the *gate closing* subprocess, we conducted an ANOVA with GATE SWITCH (switch comparison, no-switch comparison) as the within-subject factor and AGE GROUP (younger, older) as the between-subject factor. A significant main effect of GATE SWITCH means a presence of *gate closing*, while a significant GATE SWITCH × AGE GROUP interaction suggests a difference in the subprocess between the age groups.

While the ANOVA analyses can show the presence of a subprocess and a difference between the age groups, and the *post hoc* tests can indicate the differences between the different conditions, a more direct comparison of the size of the effects would be to compare the difference in mean RT between the conditions. The RT differences were computed according to Table 1 and then compared between the age groups.

A case could be made that instead of the simple RT differences, relative RT differences should be analysed between the groups as the older group is relatively slower overall compared to the younger group. This argument holds if the investigated process also slows down with aging, which is a possibility. Therefore, we also calculated a RT ratio index for each participant, which was the RT difference between the conditions divided by the average RT for the conditions (the sum divided by two). In the case of *substitution*, two ratios were calculated (reference and comparison), and then subtracted.

Similar analyses were performed separately for incorrect responses and misses. The results aligned with the RT results and a summary can be found in the Supplementary Material.

#### 2.4.3. ERP preprocessing and data analysis

The EEG data was filtered offline with a non-causal Kaiser-windowed Finite Impulse Response filter (low pass filter parameters: 30 Hz cut off frequency, beta of 12.2653, a transition bandwidth of 10 Hz; high pass filter parameters: 0.1 Hz cut off frequency, beta of 5.6533, a transition bandwidth of 0.2 Hz). Independent Component Analysis (ICA) was applied on the filtered EEG data in order to reject eye-movement artifacts (blinking, horizontal eye movements). The EEG was then segmented into epochs of 1,100 ms from 100 ms pre-stimulus to 1,000 ms post-stimulus. The mean voltage during the 100 ms pre-stimulus interval served as the baseline for amplitude measurements. Stimulus onset was measured by a photodiode, providing exact zero value for averaging. Epochs were rejected if they had a larger than 100 μV voltage change between the minimum and maximum of the epoch on any channel.

Epochs for trials with correct responses were averaged per participant for each condition relevant to the calculation of the effects, and then difference potentials were computed according to Table 1: 1) *updating* = no-switch reference ERP – minus – no-switch comparison ERP, 2) *substitution* = (different no-switch reference ERP – minus – same no-switch reference ERP) – minus – (different no-switch comparison ERP – minus – same no-switch comparison ERP), 3) *gate opening* = switch reference ERP – minus – no-switch reference ERP, 4) *gate closing* = switch comparison ERP – minus – no-switch comparison ERP. Difference potentials were used for the following reasons: 1) latent components that could be present across more than one ERP component are better identified [11], 2) reduced number of factors in the ANOVA analysis which reduces the possibility for meaningless interactions and chance results [20], 3) better suited for an exploratory cluster-based permutation analysis (see below). The average number of epochs per condition for each participant can be found in Table 2.

**Table 2.** Mean number of epochs per condition per participant in each age group. Only correct trials were included in the average. Standard deviation is indicated in parentheses.

| Conditions |  |  | AGE GROUP |  |
| --- | --- | --- | --- | --- |
| TRIAL TYPE | GATE SWITCH | RESPONSE | Younger group | Older group |
| Reference | No-switch | Same | 104.08 (16.62) | 113.17 (10.49) |
|  |  | Different | 93.36 (21.00) | 85.96 (17.35) |
|  | Switch | Same | 40.71 (7.48) | 43 (5.56) |
|  |  | Different | 31.79 (6.29) | 25.26 (7.00) |
| Comparison | No-switch | Same | 100.17 (19.82) | 100.57 (12.93) |
|  |  | Different | 103.92 (20.28) | 107.61 (14.36) |
|  | Switch | Same | 38.71 (5.87) | 39 (4.57) |
|  |  | Different | 40.29 (8.05) | 40.52 (5.22) |

First, a cluster-based permutation *t*-test analysis was conducted on each difference potential within the groups for electrode sites F3, Fz, F4, C3, Cz, C4, P3, Pz, P4, O1, Oz, O2 and for all time points after stimulus onset (0 – 1,000 ms). The threshold *p*-value was set at .05, and the familywise error (FWER) correction was set at .05. We used *t_max_* statistics. For each analysis, 10,000 permutations were run, and the tests were two-tailed to identify both positive and negative clusters. The cluster-based permutation analysis was chosen as an exploratory approach as to when (e.g. early or late processes) and where (e.g. anteriorly or posteriorly) the investigated effect was present (if at all) in each age group. The full results are presented in the Supplementary Material.

Next, we identified time windows in which to compare the difference potentials between the age groups. The time windows were selected based on the results from the cluster-based permutation analysis and visual inspection of the difference potentials (see Results). For *updating*, the selected time windows were 80-180 ms, 300-400 ms, and 400-1,000 ms post-stimulus. For *substitution*, the selected time windows were 250-600 ms and 550-750 ms post-stimulus. For *gate opening*, the selected time windows were 150-250 ms, 300-500 ms, and 500- 700 ms post-stimulus. For *gate closing*, the selected time windows were 150-300 ms and 300- 600 ms post-stimulus. Mixed ANOVAs were performed for each condition and each time window across the midline with ANTERIORITY (Fz, Cz, Pz, Oz) as the within-subject factor and AGE GROUP (younger, older) as the between-subject factor.

EEG data were preprocessed with MATLAB R2015a [21] and EEGLAB [22]. ICA was performed with the runica function of the ERPLAB toolbox [23]. A cluster-based permutation *t*-test analysis was conducted on each difference potential within the groups with the Mass Univariate ERP Toolbox [24].

#### 2.4.4. sLORETA analysis

We applied a distributed source localisation technique to locate and compare the cortical sources for potential differences between the groups. The source signal of the average ERP time series was reconstructed on the cortical surface by applying the sLORETA inverse solution [25]. The sLORETA gives a solution for the EEG inverse problem by applying a weighted minimum norm estimation with spatial smoothing and standardization of the current density map. The forward model was generated on a realistic BEM head model [26] by applying a template MRI (ICBM152; 1 mm3 voxel resolution) with template electrode positions. The reconstructed dipoles (pA/m) were determined for every 15,002 sources in three orthogonal directions (unconstrained solution). For each subject and each subprocess, the sources for the respective ERPs (see ERP preprocessing and data analysis) were estimated, their difference computed, then normalized to baseline and flattened.

For the comparison between the groups non-parametric two-sample independent t-tests were conducted on the source differences for the same time intervals as defined in the difference potential analysis, and FDR correction (alpha = .01) was applied [27]. Only differences in at least five voxels were considered. Brain regions for the corresponding significant activations were identified based on the parcellation scheme introduced by Klein and Tourville [28]. A source localisation analysis regarding the sources of the effects within each group can be found in the Supplementary Material.

The sLORETA analysis was performed with Brainstorm [29]. Group analysis was conducted according to the Group analysis processing pipeline described in Tadel et al. [27].

#### 2.4.5. Bayesian analysis

There was no *a priori* power analysis to calculate the size of the sample, thus Bayesian analyses were also conducted to evaluate the strength of the evidence for either the null or the alternative hypothesis [30,31]. The default prior distributions for ANOVA in JASP were used. Specifically, fixed effects had an *r* scale of 0.5, random effects had an *r* scale of 1, and covariates had an *r* scale of 0.354 (where covariates were included in analyses). Factors were compared across matched models. We also used the default prior option for the *t*-tests and the Mann-Whitney nonparametric tests, a Cauchy distribution with spread *r* set to 0.707. All tests were two-tailed. A Bayesian Factor (BF) larger than 3 indicates evidence for the alternative hypothesis, while a BF smaller than 0.333 suggests evidence for the null hypothesis.

The statistical analyses including the Bayesian analysis were performed with JASP [32]. The Greenhouse-Geisser corrections were used when necessary. For *post-hoc* comparisons the Bonferroni correction was applied. For the comparisons of the size of the effects between the groups an independent *t*-test or a Mann-Whitney nonparametric test was applied depending on the assumption of normality.

## 3. Results

The mean RTs for each condition and age group can be found in Table 3. The mean amplitudes for each difference potential, time window, and age group are displayed in Table 4. The relevant statistical behavioural and ERP results are summarised in **Error! Reference source not found.** and Table 6, respectively. The results from the source localisation analyses are summarised in Table 7. Full details regarding the descriptive statistics and the statistical results can be found in the Supplementary Tables 1, 2, and 3.

**Table 3.** Mean reaction time for correct trials per condition per participant in each age group. Standard deviation is indicated in parentheses.

| Conditions |  |  | AGE GROUP |  |
| --- | --- | --- | --- | --- |
| TRIAL TYPE | GATE SWITCH | RESPONSE | Younger group | Older group |
| Overall |  |  | 521.76 (97.01) | 652.41 (118.5) |
| Reference | No-switch | Same | 476.76 (91.22) | 577.61 (113.75) |
|  |  | Different | 591.57 (109.49) | 775.32 (148.79) |
|  | Switch | Same | 508.87 (125.89) | 616.38 (128.51) |
|  |  | Different | 625.23 (147.91) | 846.95 (170.7) |
| Comparison | No-switch | Same | 451.32 (82.91) | 578.25 (106.35) |
|  |  | Different | 510.72 (83.95) | 652.72 (114.83) |
|  | Switch | Same | 526.05 (120.78) | 613.73 (122.36) |
|  |  | Different | 620 (157.48) | 742.3 (140.36) |

**Table 4.** Mean amplitude per effect for each time window and each electrode site in each age group. Standard deviation is indicated in parentheses.

| Age group |  | Younger group |  |  |  | Older group |  |  |  |
| --- | --- | --- | --- | --- | --- | --- | --- | --- | --- |
| Electrode site |  | FZ | CZ | PZ | OZ | FZ | CZ | PZ | OZ |
| Effect | Time window |  |  |  |  |  |  |  |  |
| Updating | 80-180 ms | 0.37<br>(0.73) | -0.42<br>(0.74) | -0.88<br>(1.79) | -1.91<br>(2.22) | 0.17<br>(0.47) | -0.198<br>(0.44) | -0.44<br>(0.87) | -0.996<br>(1.47) |
|  | 300-400 ms | 0.01<br>(0.4) | -0.15<br>(0.49) | -0.15<br>(0.9) | -1.995<br>(1.39) | 0.12<br>(0.32) | -0.07<br>(0.29) | -0.08<br>(0.56) | -0.75<br>(0.85) |
|  | 400-1,000 ms | 0.08<br>(0.29) | -0.14<br>(0.26) | -0.62<br>(0.55) | -1.795<br>(1.15) | -0.12<br>(0.24) | -0.02<br>(0.21) | -0.26<br>(0.51) | -1.67<br>(0.84) |
| Substitution | 250-500 ms | 0.77<br>(0.78) | -0.29<br>(0.98) | -0.58<br>(1.73) | 0.39<br>(2.18) | -0.08<br>(0.73) | 0.24<br>(0.53) | 0.34<br>(1.38) | 0.15<br>(1.69) |
|  | 500-600 ms | 0.85<br>(1.24) | -0.25<br>(1.19) | 0.05<br>(2.42) | 0.59<br>(3.24) | 0.43<br>(0.87) | -0.15<br>(0.92) | 0.82<br>(2.07) | 1.30<br>(2.37) |
|  | 650-750 ms | 0.08<br>(0.89) | -0.00<br>(1.05) | -0.12<br>(1.91) | -0.54<br>(2.51) | 0.34<br>(0.59) | -0.03<br>(0.74) | 0.96<br>(1.77) | 1.5<br>(2.15) |
| Gate opening | 150-250 ms | -0.3<br>(0.48) | 0.27<br>(0.74) | 0.43<br>(1.48) | -0.21<br>(1.7) | -0.06<br>(0.69) | -0.28<br>(0.51) | -0.68<br>(1.14) | -1.66<br>(1.41) |
|  | 300-500 ms | -0.44<br>(0.66) | 0.76<br>(0.9) | 1.86<br>(2.27) | 1.45<br>(2.82) | 0.01<br>(0.56) | 0.04<br>(0.52) | 0.32<br>(1.17) | -0.08<br>(1.53) |
|  | 500-700 ms | -0.33<br>(0.68) | 0.64<br>(0.78) | 1.4<br>(1.88) | 1.46<br>(2.65) | 0.2<br>(0.52) | 0.06<br>(0.5) | 0.56<br>(1.08) | 0.75<br>(1.16) |
| Gate closing | 150-300 ms | 0.7<br>(0.56) | -0.16<br>(0.55) | -0.42<br>(0.93) | -0.92<br>(1.28) | -0.09<br>(0.34) | -0.12<br>(0.44) | -0.37<br>(0.99) | -0.91<br>(1.15) |
|  | 300-600 ms | 0.42<br>(0.62) | -0.03<br>(0.57) | 0.26<br>(1.57) | -0.36<br>(1.8) | 0.32<br>(0.52) | -0.34<br>(0.45) | -0.12<br>(0.96) | 0.05<br>(1.44) |

**Table 5.**
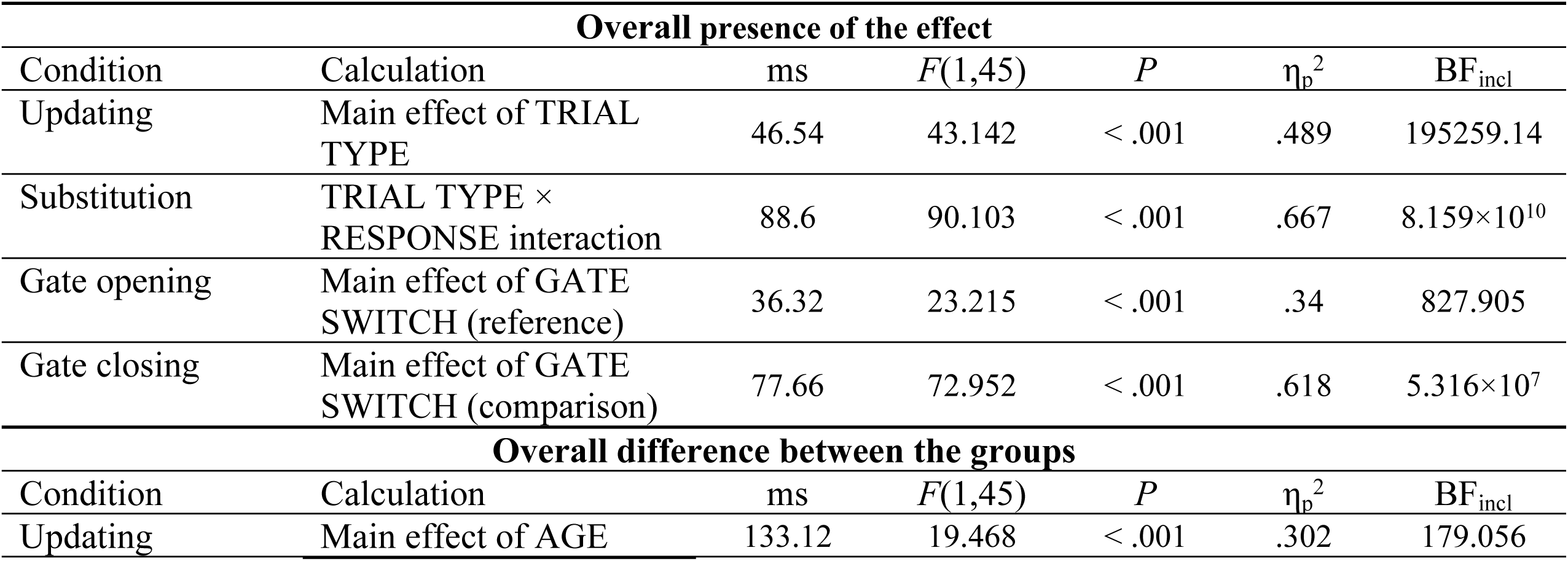
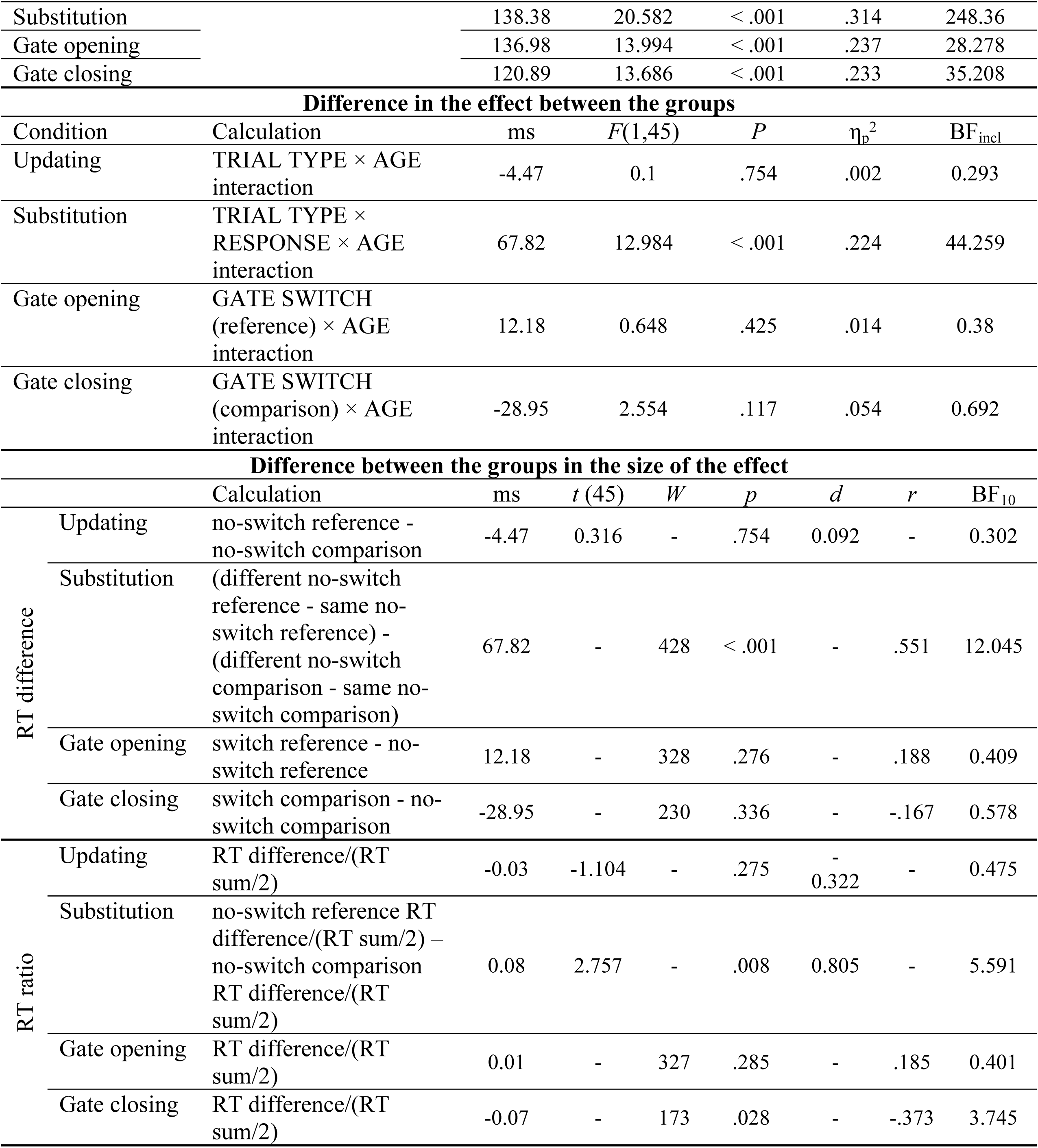
A summary of the behavioural statistical results regarding the overall presence of the subprocess, the differences in the subprocess between the groups (repeated measures ANOVA results), and the differences in the size of the effects (independent groups comparisons). The ms column indicates the RT difference.

**Table 6.** A summary of the ERP statistical results regarding the differences in the subprocesses between the groups and between electrode sites.

| **Difference in the effect between the groups (Main effect of AGE)** |  |  |  |  |  |  |
| --- | --- | --- | --- | --- | --- | --- |
| Condition | Time Window | $F(1,45)$ | $P$ | $\eta_p^2$ | $\epsilon$ | $BF_{incl}$ |
| Updating | 80-180 ms | 0.756 | .389 | .017 | - | 0.312 |
|  | 300-400 ms | 1.609 | .211 | .035 | - | 0.522 |
|  | 400-1,000 ms | 5.818 | .02 | .114 | - | 2.541 |
| Substitution | 250-500 ms | 0.098 | 0.755 | 0.002 | - | 0.254 |
|  | 500-600 ms | 0.438 | 0.511 | 0.01 | - | 0.344 |
|  | 650-750 ms | 5.838 | .02 | .115 | - | 2.673 |
| Gate opening | 150-250 ms | 9.168 | .004 | .169 | - | 9.413 |
|  | 300-500 ms | 5.675 | .021 | .112 | - | 2.778 |
|  | 500-700 ms | 1.877 | .177 | .04 | - | 0.579 |
| Gate closing | 150-300 ms | 0.058 | .81 | .001 | - | 0.307 |
|  | 300-600 ms | 0.153 | .696 | .003 | - | 0.286 |
| **Difference in the effect between electrode sites (Main effect of ANTERIORITY)** |  |  |  |  |  |  |
| Condition | Time Window | $F(1,45)$ | $P$ | $\eta_p^2$ | $\epsilon$ | $BF_{incl}$ |
| Updating | 80-180 ms | 103.262 | < .001 | .696 | .536 | $2.055 \times 10^{34}$ |
| | 300-400 ms | 22.762 | < .001 | .336 | .608 | $1.714 \times 10^9$ |
| | 400-1,000 ms | 53.639 | < .001 | .544 | .639 | $1.549 \times 10^{18}$ |
| Substitution | 250-500 ms | 1.756 | 0.174 | 0.038 | 0.743 | 0.221 |
|  | 500-600 ms | 4.594 | 0.01 | 0.093 | 0.736 | 6.115 |
|  | 650-750 ms | 1.413 | .242 | .03 | .686 | 0.113 |
| Gate opening | 150-250 ms | 9.785 | < .001 | .179 | .573 | 610.869 |
|  | 300-500 ms | 10.337 | < .001 | .187 | .48 | 1757.509 |
|  | 500-700 ms | 11.4 | < .001 | .202 | .611 | 11554.696 |
| Gate closing | 150-300 ms | 19.183 | < .001 | .299 | .676 | $7.503 \times 10^7$ |
|  | 300-600 ms | 3.85 | .027 | .079 | .633 | 2.729 |
| **Difference in the effect between the groups depending on electrode site (AGE × ANTERIORITY interaction)** |  |  |  |  |  |  |
| Condition | Time Window | $F(1,45)$ | $P$ | $\eta_p^2$ | $\epsilon$ | $BF_{incl}$ |
| Updating | 80-180 ms | 2.079 | .142 | .044 | .536 | 0.573 |
|  | 300-400 ms | 2.36 | .106 | .05 | .608 | 0.764 |
|  | 400-1,000 ms | 10.375 | < .001 | .187 | .639 | 5485.974 |
| Substitution | 250-500 ms | 5.473 | 0.004 | 0.108 | 0.743 | 29.109 |
|  | 500-600 ms | 1.563 | 0.213 | 0.034 | 0.736 | 0.316 |
|  | 650-750 ms | 6.063 | .003 | .119 | .686 | 51.42 |
| Gate opening | 150-250 ms | 7.589 | .002 | .144 | .573 | 257.032 |
|  | 300-500 ms | 7.568 | .003 | .144 | .48 | 253.923 |
|  | 500-700 ms | 3.73 | .032 | .077 | .611 | 3.577 |
| Gate closing | 150-300 ms | 0.699 | .502 | .015 | .676 | 0.122 |
|  | 300-600 ms | 1.809 | .172 | .039 | .633 | 0.405 |

**Table 7.** Source differences between the younger and the older group. Regions, left (L) or right (R) hemisphere, number of significant voxels in the region as well as the MNI (x,y,z), t-value, and p-value of the peak difference voxel are shown.

| Region | HM | $N_{voxel}$ | MNI coordinates | | | $t$ | $p$ |
| --- | --- | --- | --- | --- | --- | --- | --- |
|  |  |  | X | y | z |  |  |
| *Updating* |  |  |  |  |  |  |  |
| 400-1000 ms |  |  |  |  |  |  |  |
| Lateral orbitofrontal gyrus | R | 21 | 30.9 | 20.9 | -26.2 | 3.786 | 0.0002 |
| Insula | R | 19 | 26.8 | 7.4 | -25.7 | 3.639 | 0.0002 |
| Inferior temporal gyrus | R | 42 | 46.8 | 1.8 | -48.3 | 4.235 | 0.0002 |
| Middle temporal gyrus | R | 47 | 50.6 | 4.9 | -43.0 | 4.210 | 0.0002 |
| Superior temporal gyrus | R | 52 | 47.9 | 22.1 | -27.5 | 4.169 | 0.0002 |
| Temporal pole | R | 33 | 37.8 | 25.0 | -36.3 | 4.009 | 0.0002 |
| Entorhinal cortex | R | 28 | 27.0 | -0.8 | -39.5 | 3.709 | 0.0002 |
| Fusiform gyrus | R | 22 | 29.2 | -3.7 | -51.7 | 3.812 | 0.0002 |
| *Gate opening* |  |  |  |  |  |  |  |
| 500-700 ms |  |  |  |  |  |  |  |
| Rostral anterior cingulate | L | 21 | -1.4 | 14.2 | -14.7 | 3.375 | 0.0004 |
|  | R | 6 | 2.1 | 17.4 | -12.1 | 3.296 | 0.0006 |
| Lateral orbitofrontal gyrus | L | 108 | -21.1 | 9.9 | -27.8 | 3.640 | 0.0004 |
| Medial orbitofrontal gyrus | L | 113 | -14.8 | 7.6 | -21.9 | 3.537 | 0.0004 |
|  | R | 42 | 4.7 | 8.1 | -20.2 | 3.549 | 0.0010 |
| Precentral gyrus | L | 9 | -43.4 | 0.4 | 5.9 | 3.375 | 0.0008 |
| Insula | L | 83 | -25.8 | 6.2 | -21.0 | 3.607 | 0.0004 |
|  | R | 7 | 2.1 | 7.8 | -15.6 | 3.535 | 0.0008 |
| Lingual gyrus | L | 12 | -15.6 | -43.4 | -14.2 | 3.643 | 0.0008 |
| Inferior temporal gyrus | L | 129 | -30.5 | 8.5 | -41.2 | 3.549 | 0.0004 |
| Middle temporal gyrus | L | 119 | -53.6 | -20.1 | -14.0 | 3.532 | 0.0002 |
| Superior temporal gyrus | L | 111 | -32.7 | 10.5 | -26.0 | 3.461 | 0.0004 |
| Temporal pole | L | 27 | -21.9 | 9.9 | -32.0 | 3.627 | 0.0004 |
| Transverse temporal gyrus | L | 11 | -33.0 | -25.3 | 8.1 | 3.421 | 0.0002 |
| Entorhinal cortex | L | 27 | -15.5 | -16.7 | -25.7 | 3.932 | 0.0004 |
| Fusiform gyrus | L | 113 | -21.6 | -38.3 | -18.7 | 3.737 | 0.0006 |
| Parahippocampal gyrus | L | 61 | -16.5 | -25.0 | -19.4 | 4.136 | 0.0004 |

### 3.1. Overall behavioural results

Overall accuracy was relatively high in both groups (*M*=85.9% in the younger and *M*=80.8% in the older group). The accuracy was somewhat lower in the older group than in the younger group, *U* = 154, *p* = .01, *r* = -0.442, BF_10_ = 4.858. This was due to the older group having a higher rate of misses than the younger group (5% vs. 3.1%); *U* = 422, *p* = .002, *r* = 0.529, BF_10_ = 9.387; while the evidence for a higher rate of incorrect responses in the older group was insufficient, *U* = 373.5, *p* = .039, *r* = 0.353, BF_10_ = 1.477 (11% in the younger vs. 14.1% in the older group).

The older participants were slower overall than the younger group (521.76 ms in the younger vs. 652.41 ms in the older group), *U* = 434.00, *p* < .001, *r* = 0.572, BF_10_ = 29.783.

### 3.2. Updating

Figure 2 displays the behavioural and the ERP results.

**Figure 2.**
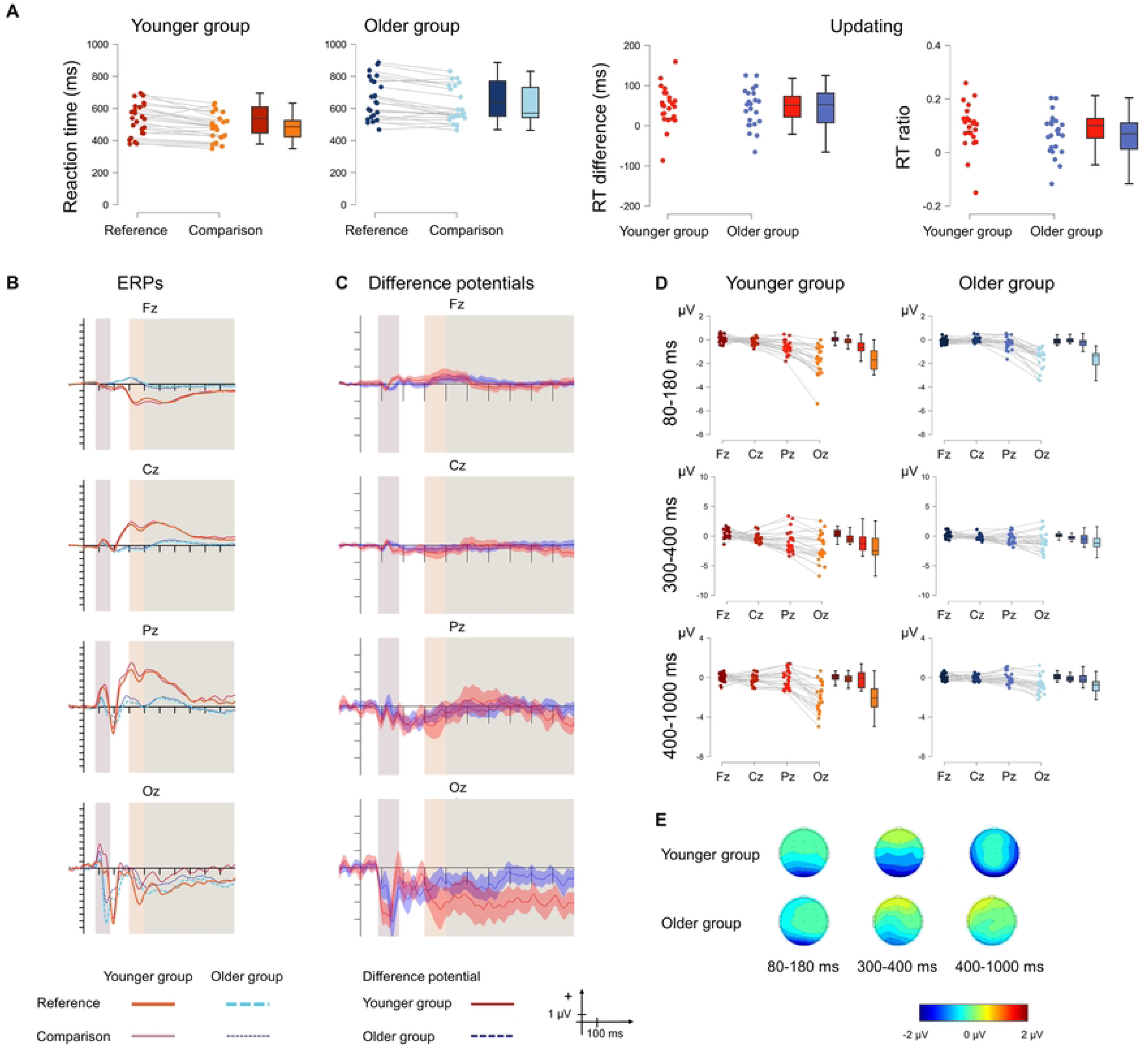
**A** Behavioural statistical results for *updating* in the younger (red) and the older (blue) groups as well as for the size of the effect (right panel). The darker shade shows the reference condition, and the lighter shade shows the comparison condition. **B** ERPs to the no-switch reference and no-switch comparison conditions and **C** Difference potentials with 95% CI intervals. Red continuous lines indicate the younger group, while blue dashed lines indicate the older group. For the ERPs the thick lines show the no-switch reference ERPs and the thin lines show the no-switch comparison ERPs. The shaded areas show the three time windows for analysis. **D** ERP statistical results for the difference in the updating effect between the two age groups for the 80-180 ms (top), 300-400 ms (middle) and 400-1,000 ms (bottom) time windows. Red colour indicates the younger group and blue colour indicates the older group. **E** Scalp distributions for the 80-180 ms (left), 300-400 ms (middle) and 400-1,000 ms (right) time windows for the younger (top) and the older (bottom) groups.

#### 3.2.1. Behavioural results

As expected, the older participants were slower overall than the younger participants (639 ms vs. 506 ms). The *updating cost* was present (594 ms for reference vs. 548 ms for comparison trials), but there was no difference in the effect between the groups with moderate evidence for the null hypothesis. A comparison of the size of the effect between the two age groups confirmed that there was no difference either for the RT difference (48.72 ms in the younger and 44.25 ms in the older group) or the RT ratio (0.093 in the younger and 0.067 in the older group).

#### 3.2.2. ERP results

In the 80-180 ms time window, there was moderate evidence for no main effect of AGE GROUP. The main effect of ANTERIORITY was significant. The amplitude measured at Oz was significantly more negative than those measured at Fz, Cz, and Pz (all *p* < .001). The amplitude measured at Pz was significantly more negative from those measured at both Fz and Cz (*p* = .002, and *p* = .009, respectively). The interaction was not significant but the evidence for the lack of effect was insufficient.

In the 300-400 ms time window, there was insufficient evidence for a main effect of AGE GROUP. The main effect of ANTERIORITY was significant. The amplitude measured at Fz was significantly more positive than that measured at Cz, Pz, and Oz (all *p* < .001). The amplitude measured at Oz was significantly more negative than those measured at Cz and Pz (*p* < .001 and *p* = .002, respectively). The evidence for an interaction was insufficient.

In the 400-1,000 ms time window, there was only anecdotal evidence for a main effect of AGE GROUP. The main effect of ANTERIORITY was significant. The amplitude measured at Oz was significantly more negative from those measured at Fz, Cz, and Pz (all *p* < .001). The interaction was also significant, and was caused by a significant difference in amplitude between the younger and the older group at Oz (*p* < .001), with the amplitude being more negative in the younger group.

#### 3.2.3. Source localisation results

Between-group differences were found only in the 400-1,000 ms time interval (Table 1 and Figure 6). Activation was larger in the younger group in the right hemisphere, in frontal (lateral orbitofrontal gyrus and insula) and temporal (inferior temporal gyrus, middle temporal gyrus, superior temporal gyrus, temporal pole, entorhinal cortex, fusiform gyrus) regions.

### 3.3. Substitution

Figure 3 displays the behavioural and the ERP results.

**Figure 3.**
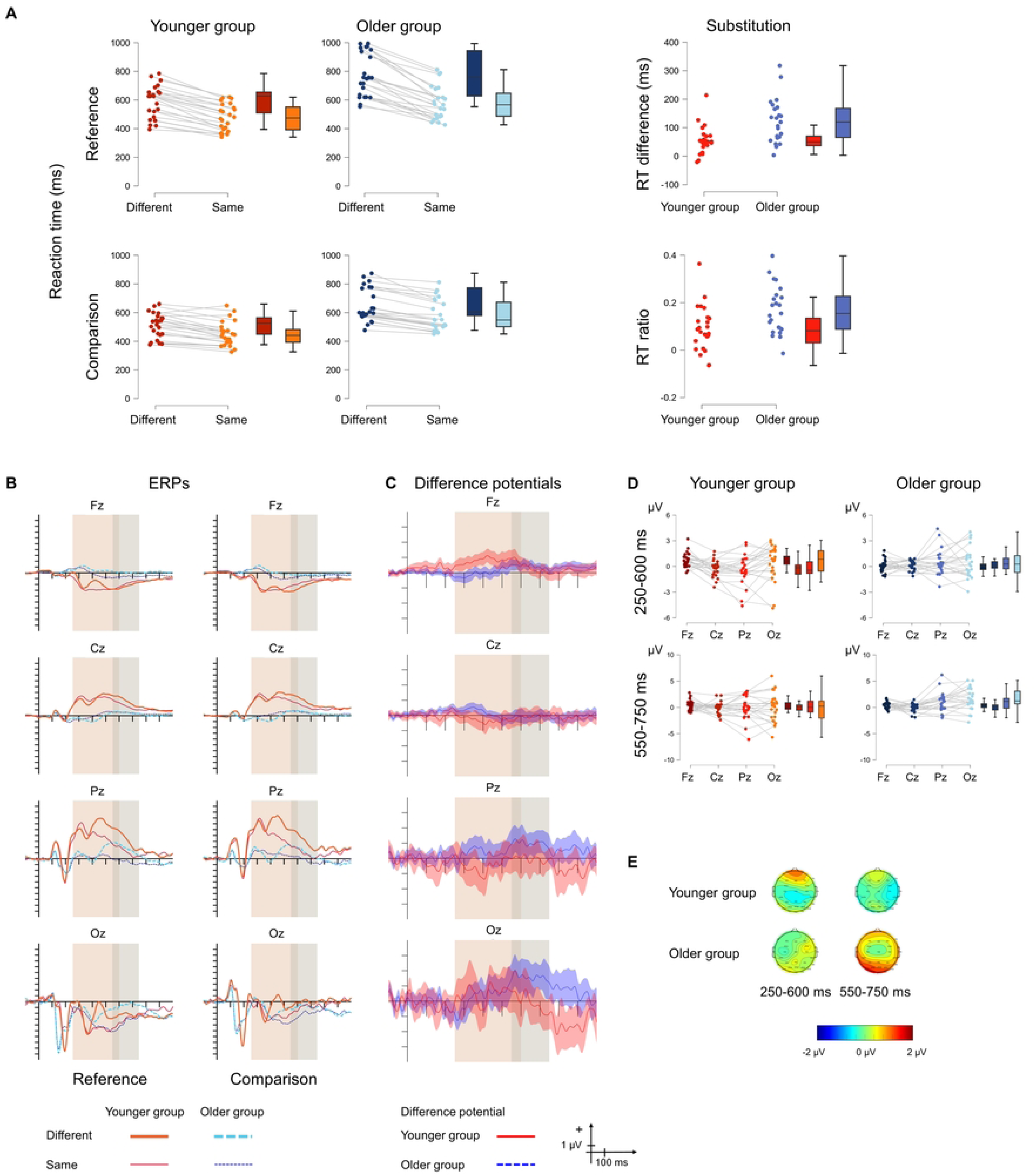
**A** Behavioural statistical results for *substitution* in the younger (red) and the older (blue) groups as well as for the size of the effect (right panel). The two panels on the left show the no-switch reference trials, and the two panels on the right show the no-switch comparison trials. The darker shade shows the “different”, and the lighter shade shows the “same” condition. **B** ERPs to the no-switch reference (left) and no-switch comparison (right) conditions and **C** Difference potentials with 95% CI intervals. Red continuous lines indicate the younger group, while blue dashed lines indicate the older group. For the ERPs the thick lines show the “different” ERPs, and the thin lines show the “same” ERPs. The shaded areas show the two time windows for analysis. **D** ERP statistical results for the difference in the substitution effect between the two age groups for the 250-600 ms (top) and 550-750 ms (bottom) time windows. Red colour indicates the younger group and blue colour indicates the older group. **E** Scalp distributions for the 250-600 ms (left) and 550-750 ms (right) time windows for the younger (top) and the older (bottom) groups.

#### 3.3.1. Behavioural results

The older group was slower overall than the younger group (646 ms vs. 508 ms). *Substitution* was significant as seen in the TRIAL TYPE × RESPONSE interaction. While participants were slower overall for both different no-switch reference stimuli compared to same no-switch reference stimuli (682 ms vs. 526 ms, *p* < .001); and different no-switch comparison compared to same no-switch comparison stimuli (580 ms vs. 513 ms, *p* < .001); there was a significant difference between no-switch reference and comparison stimuli in the case of stimuli that required a “different” response (682 ms vs. 580 ms, *p* < .001); but not in the case of stimuli with the “same” response (526 ms vs. 513 ms, *p* = .721). There was also a significant difference in *substitution* between the groups as shown by the triple interaction. The older group was slower overall than the younger group, and the difference was significant in the following conditions: 1) different no-switch reference stimuli, 592 ms in the younger and 775 ms in the older group, *p* < .001; 2) different no-switch comparison stimuli, 510 ms in the younger and 653 ms in the older group, *p* = .001; and 3) same no-switch comparison stimuli, 451 ms in the younger and 578 ms in the older group, *p* = .006; but 4) the groups did not differ for the same no-switch reference stimuli, 477 ms in the younger and 577 ms in the older group, *p* = .077. A direct comparison of the size of the substitution effect showed that the effect was larger in the older than in the younger group for both RT difference (123 ms vs. 55 ms) and RT ratio (0.169 vs. 0.09).

#### 3.3.2. ERP results

In the 250-600 ms time window, there was a tendency for a main effect of ANTERIORITY, however, the evidence was inconclusive. The absence of a main effect of AGE GROUP was supported by moderate evidence for the null hypothesis. The interaction was significant and due to the mean amplitude measured at Fz being significantly more positive than that measured at Cz (.029) and Pz (*p* = .007) in the younger group.

In the 550-750 ms time window, there was a tendency for a main effect of both AGE GROUP and ANTERIORITY, but the evidence was inconclusive. The interaction was significant: there was an anteriority effect only in the older group, in which the amplitude measured at Oz was significantly more positive than that measured at Cz (*p* = .002). The younger and the older group differed significantly at the Oz electrode site (*p* = .034), with the amplitude being more positive in the older compared to the younger group.

#### 3.3.3. Source localisation results

There were no source differences between the groups.

### 3.4. Gate opening

Figure 4 displays the behavioural and the ERP results.

**Figure 4.**
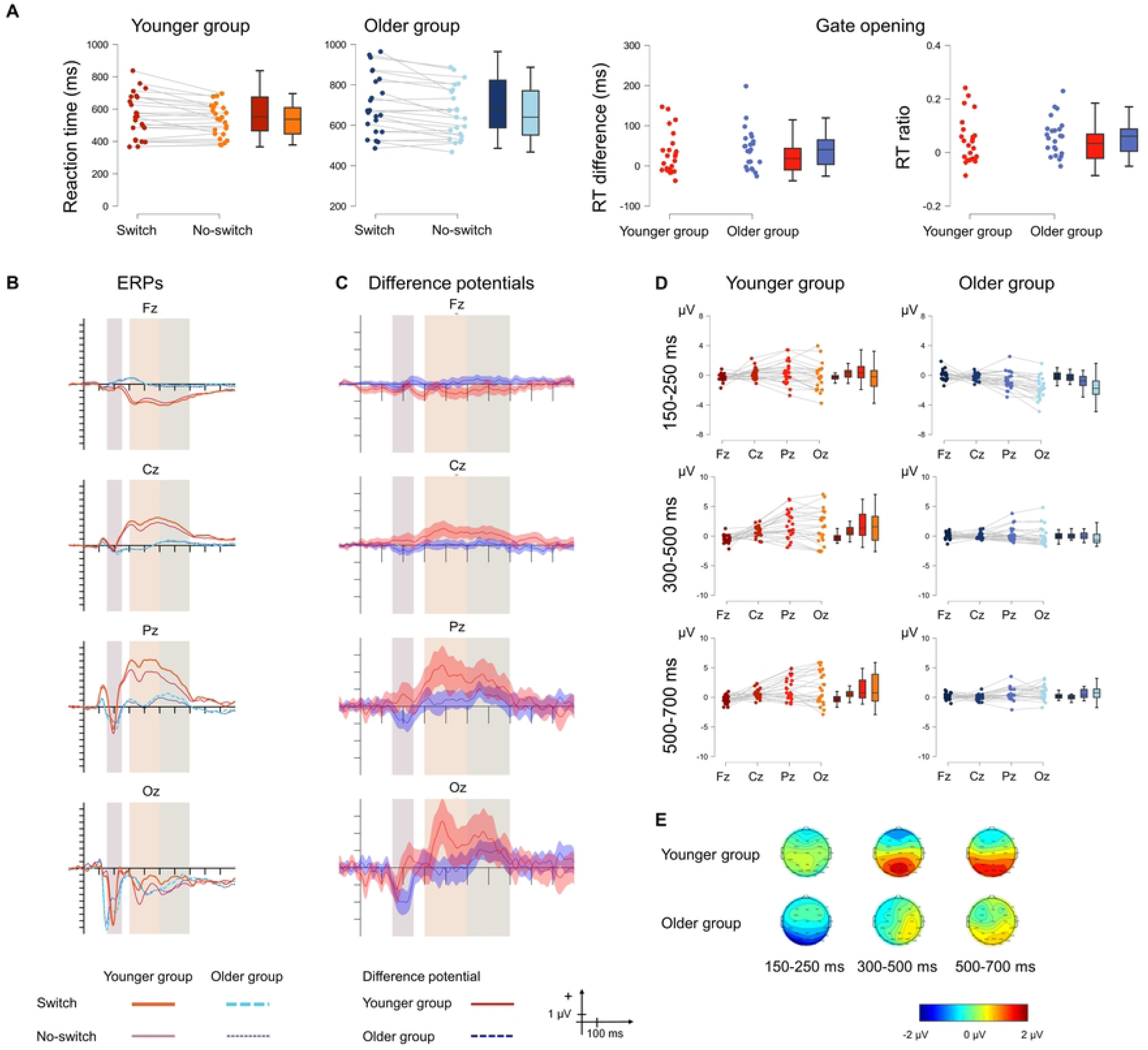
**A** Behavioural statistical results for *gate opening* in the younger (red) and the older (blue) groups as well as for the size of the effect (right panel). The darker shade shows the switch reference condition, and the lighter shade shows the no-switch reference condition. **B** ERPs to the switch reference and no-switch reference conditions and **C** Difference potentials with 95% CI intervals. Red continuous lines indicate the younger group, while blue dashed lines indicate the older group. For the ERPs the thick lines show the switch reference ERPs, and the thin lines show the no-switch reference ERPs. The shaded areas show the three time windows for analysis. **D** ERP statistical results for the difference in the gate opening effect between the two age groups for the 150-250 ms (top), 300-500 ms (middle) and 500-700 ms (bottom) time windows. Red colour indicates the younger group and blue colour indicates the older group. **E** Scalp distributions for the 150-250 ms (left), 300-500 ms (middle) and 500-700 ms (right) time windows for the younger (top) and the older (bottom) groups.

#### 3.4.1 Behavioural results

The older group was slower overall than the younger group (682 ms vs. 545 ms). *Gate opening* was indicated by switch reference trials being slower than no-switch reference trials (630 ms vs. 594 ms). The GATE SWITCH × AGE interaction was not significant with inconclusive evidence regarding the lack of difference between the groups. This result was also supported by the direct comparison of the size of the effect, which was 42.54 ms in the older and 30.36 ms in the younger group for the RT difference, and 0.059 in the older and 0.045 in the younger group for the RT ratio.

#### 3.4.2. ERP results

In the 150-250 ms time window, there was a main effect of AGE GROUP. The measured amplitude was overall more negative in the older group. The main effect of ANTERIORITY was also significant. The amplitude measured at Pz was significantly more negative than those measured at Fz and Cz (*p* < .001 and *p* = .029, respectively). The amplitude measured at Oz was significantly more negative than that measured at Fz (*p* = .002). The interaction was significant. The anteriority effect was only seen in the older group with the amplitude measured at Oz being significantly more negative than those measured at Fz, Cz, and Pz (*p* < .001, *p* < .001, and *p* = .013, respectively). The two age groups differed significantly at the Pz (*p* = .03) and Oz (*p* < .001), with the amplitude measured in the older group being more negative.

In the 300-500 ms time window, the overall amplitude in the younger group was more positive than that measured in the older group, but the evidence for a significant main effect of AGE GROUP was insufficient. The main effect of ANTERIORITY was significant. The amplitude measured at Oz was significantly more positive than those measured at Fz, Cz, and Pz (all *p* < .001). The interaction was significant. The anteriority effect was observable only in the younger group, in which the amplitudes measured at Cz, Pz, and Oz were more positive than that measured at Fz (*p* = .015, *p* < .001, and *p* < .001, respectively). The amplitude measured at Pz was more positive than that measured at Cz (*p* = .039). The amplitudes measured in the younger group at the Pz and the Cz electrode sites were more positive than those measured in the older group (*p* = .03 and *p* = .033, respectively).

In the 500-700 ms time window, there was no main effect of AGE GROUP, although the evidence for the null hypothesis was inconclusive. The main effect of ANTERIORITY was significant. The amplitude measured at Fz was significantly smaller than those measured at Pz and Oz (both *p* < .001). The amplitude measured at Cz was also significantly smaller than those measured at Pz and Oz (*p* = .042 and *p* = .008, respectively). The interaction was significant. In the younger group, the amplitude measured at Fz was significantly smaller than those measured at Pz (both *p* < .001).

#### 3.4.3 Source localisation results

There were source differences between the age groups in the 500-700 ms time interval (Table 7 and Figure 6). Activation was larger in the younger compared to the older group in the left hemisphere, mainly in frontal (orbitofrontal gyrus, precentral gyrus, insula, rostral anterior cingulate) and temporal (inferior temporal gyrus, middle temporal gyrus, superior temporal gyrus, temporal pole, transverse temporal gyrus, entorhinal cortex, fusiform gyrus, parahippocampal gyrus) regions. There was also larger activation in the younger compared to the older group in the right hemisphere, in frontal regions (medial orbitofrontal cortex, insula, rostral anterior cingulate).

### 3.5. Gate closing

Figure 5 displays the behavioural and the ERP results.

**Figure 5.**
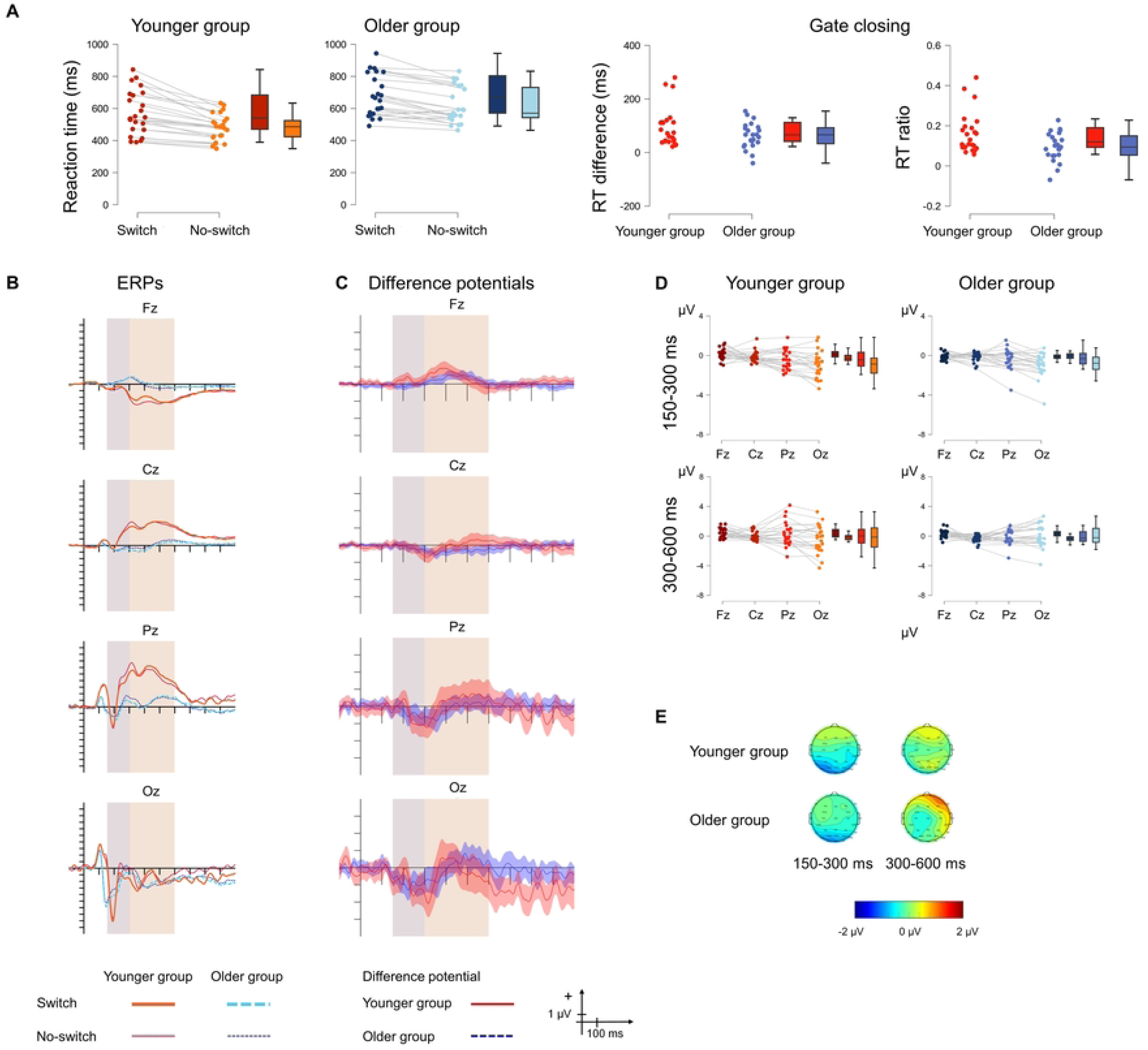
**A** Behavioural statistical results for *gate closing* in the younger (red) and the older (blue) groups as well as for the size of the effect (right panel). The darker shade shows the switch comparison condition, and the lighter shade shows the no-switch comparison condition. **B** ERPs to the switch comparison and no-switch comparison conditions and **C** Difference potentials with 95% CI intervals. Red continuous lines indicate the younger group, while blue dashed lines indicate the older group. For the ERPs the thick lines show the switch comparison ERPs, and the thin lines show the no-switch comparison ERPs. The shaded areas show the two time windows for analysis. **D** ERP statistical results for the difference in *gate closing* between the two age groups for the 150-300 ms (top) and 300-600 ms (bottom) time windows. Red colour indicates the younger group and blue colour indicates the older group. **E** Scalp distributions for the 150-300 ms (left) and 300-600 ms (right) time windows for the younger (top) and the older (bottom) groups.

**Figure 6.**
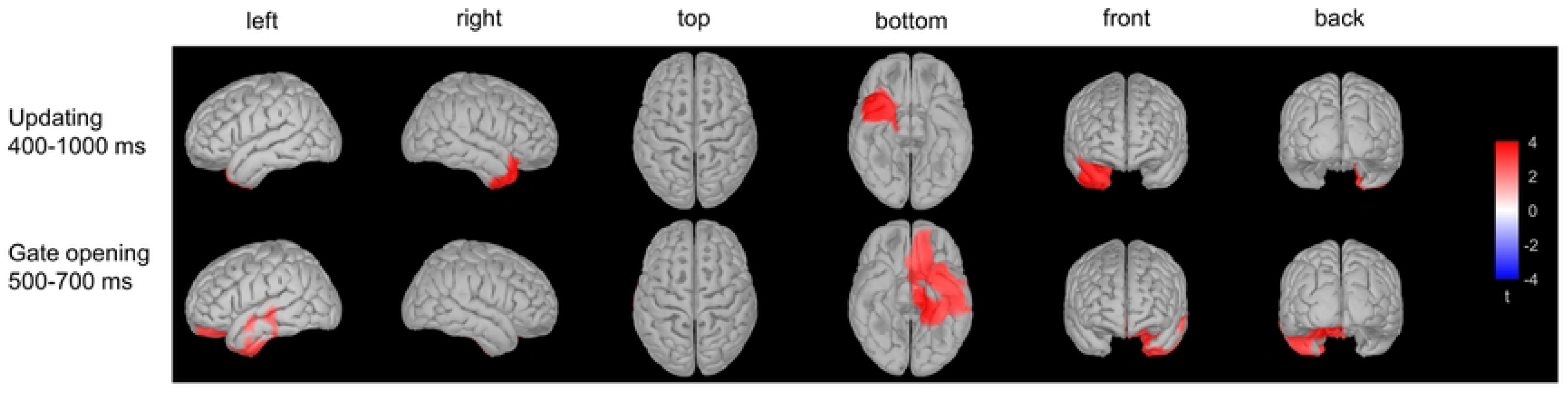
Source differences between the two age groups in *updating* (top) in the 400-1,000 ms time window and in *gate opening* (bottom) in the 500-700 ms time window.

#### 3.5.1. Behavioural results

The older group was slower overall than the younger group, 648 ms vs. 527 ms. *Gate closing* was also present overall, 625 ms for switch comparison compared to 548 ms for no- switch comparison stimuli. There was no difference between the groups with insufficient evidence for the null hypothesis. The direct comparison of the size of *gate closing* between the older (62.88 ms) and the younger (91.83 ms) also showed no significant difference for the RT difference. Regarding the RT ratio, a smaller gate closing effect is observed in the older (0.095) compared to the younger group (0.161). Of note, the difference seems to be driven by three outlier values in the younger group, and the removal of the outliers leads to only a trend in the difference, *t*(45) = -1.745, *p* = 0.088, *Cohen’s d* = -0.527, BF_10_ = 0.995.

#### 3.5.2. ERP results

In the 150-300 ms time window, there was moderate evidence for main effect of AGE GROUP. The mean amplitude was more negative in the younger than in the older group. The main effect of ANTERIORITY was significant. The amplitude measured at Oz was significantly more negative than those measured at Fz, Cz, and Pz (*p* < .001, *p* < .001, and *p* = .001, respectively). The amplitude measured at Pz was significantly more negative than that measured at Fz (*p* = .007). There was moderate evidence for a lack of interaction.

In the 300-600 ms time window, there was moderate evidence for no main effect of AGE GROUP. There was insufficient evidence for a main effect of ANTERIORITY. There was also insufficient evidence for a lack of interaction.

#### 3.5.3. Source localisation results

There were no source differences between the groups.

## 4. Discussion

Our study employed a data-based approach to explore the electrophysiological correlates of the working memory subprocesses revealed by the reference-back paradigm. Given the absence of prior results, we specifically investigated which subprocesses are influenced by aging, and how.

We replicated the experimental design of the reference-back paradigm initially introduced by Rac-Lubashevsky and Kessler [2] and obtained similar behavioural results. We observed significant costs for all four subprocesses we measured: *updating*, *substitution*, *gate opening* and *gate closing*. These results, consistent with others [3,9], confirmed the robustness of these subprocesses. Beyond the usual age-related slowing in reaction time, we found differences between the age groups only in *substitution*. This cost of updating working memory with new information was larger in older compared to younger adults. As the RT ratio method eliminates age-related slowing, we might conclude from behavioural results that *substitution* is the most vulnerable among the examined subprocesses in the elderly. However, the ERP data provides a deeper insight into what may underlie these subprocesses.

For *updating*, the cluster-based permutation analysis and visual inspection suggested three negative ERP components in both groups, which showed the processing difference between reference and comparison trials in the no-switch trials. Although we could say that this pattern in itself characterizes the *updating* process, it is worth examining what other processes may contribute to its development. Can we assume there are differences between the two trial types other than the *updating* process, which are expressed in individual components and do not necessarily characterize *updating*?

The negativity between 80-180 ms had an occipital maximum. It most likely indicates differences in visual processing and not *updating* per se. Firstly, the stimuli were physically different: while the frame colour was red in the reference trials, it was blue in the comparison trials. Secondly, the red frame was a prompt for working memory updating, and attentional resources may have been better allocated to it. This could have been indicated by the occipital N1 component which reflects a discrimination effect in the attended area [33,34]. The two age groups did not show differences in this processing stage: neither in the size of the amplitude, nor in its distribution, nor in the source of the component.

The negativity observed between 300-400 ms also showed an occipital maximum, and demonstrated anteriority, but did not manifest any other significant effects. The component may belong to the N2 family and can reflect selective attention [35]. Given that the reference and comparison trials have different significance in the task, it is reasonable to assume that the attention given to them is not entirely the same, as the greater significance of the reference trials was reflected in its larger amplitude.

Typically, within this time window, we would expect the P3b component to characterize the updating of working memory content, resulting in a positive component seen parietally [10,36]. Using the reference-back paradigm, Rac-Lubashevsky and Kessler [8] could not confirm that there would be a connection between P3b and WM updating; instead, they concluded that P3b reflected target categorization process. In contrast to Rac-Lubashevsky and Kessler, we analysed the difference curves, isolating the neural processes that differ in the two trial types; and consequently, characterize the *updating* process. In the older age group, the original curve lacks a prominent P3b, consistent with the idea that this component may not necessarily emerge in more challenging tasks for them [37]. Therefore, it is not surprising that the difference curve does not reveal this component in their case either. While we observed the P3b component in the original curves for younger participants, it was absent in the difference curve. Accordingly, we could conclude that the P3b component does not play a role in updating working memory content aligning with the notion that the component may indicate other processes, such as the closure of cognitive epochs [38,39]; stimulus-response link reactivation [40,41]; response facilitation [42]; or decision making [43,44]. However, whereas in Rac-Lubashevsky and Kessler’s model *updating cost* entails the updating of working memory content with both new and old information, other studies define updating as the incorporation of new information, akin to *substitution* in the current model. This implies that we should evaluate the theory that P3b reflects working memory updating when discussing the *substitution* results.

Finally, we found a significant negative shift in the younger but not the older group between 400 and 1,000 ms occipitally. In the two age groups, there are differences again in the course of the original curves: in older adults, there is hardly any difference between the two trial types, indicating less differentiation between the tasks in this later stage; whereas in younger adults, the curve returns to baseline at about 600 ms in comparison trials, while in reference trials, when updating is required, there is a noticeable large negativity during the whole epoch. Source analysis also confirmed age-group differences, as larger activity was found in the right frontal and temporal areas in the younger compared to the older adults. Similar late posterior negativity was found when the memory task required the binding of items with contextual information specifying the given episode [45].

In now summarizing the *updating* results, we suggest that the occipital negativity between 300 and 1,000 ms characterizes this process in younger adults, while it appears that older adults do not distinctly differentiate between trials that require both matching and updating, and trials that require only matching.

While the *updating cost* encompasses both the updating of working memory with new and old information, *substitution* represents a more specific process, indicating solely the updating of working memory with new information. This is calculated through a double subtraction, where we contrast the matching components within the no-switch trials between reference and comparison trials. Cluster analysis showed two significant time windows after this subtraction: a component between 250-600 ms for the younger group, and another one between 550-750 ms for the older adults. In younger adults, the positivity between 250-600 ms showed a frontal maximum, while in older adults, the positivity between 550-750 ms had a posterior maximum. Overall, in both groups, the electrophysiological signal of updating with new information differed from the pattern observed during *updating*, and these components can be compared with the P3b obtained in other experiments. Although the P3b component typically shows a later latency in the older group; its distribution is parietal in younger individuals, whereas in older adults, it may show a more uniform distribution, with a smaller parietal peak and the emergence of a frontal maximum as well [e.g., 46,47]. However, the pattern observed here suggests different underlying processes between the two age groups, which is further supported by the sLORETA results. While the group differences of the sources were not significant, the group results, shown in the Supplementary Material though not discussed in detail here, indicated a greater involvement of frontal sources in younger adults and posterior sources in older adults. Our within-group sLORETA analysis highlighted areas showing higher activity difference between the conditions relative to the average activity difference across the entire scalp. However, the within-group brain activity differences could still be similar across groups, e.g., the larger difference in the parietal region could be similar magnitude in the two groups, but of different magnitude when compared to other regions within the group. Therefore, it is possible that although different sources are found when analysing age groups separately, we do not obtain significant results when comparing groups in a whole-brain analysis. Despite this methodological limitation, this result suggests a significant difference between the two age groups in one of the subprocesses of working memory as different brain regions are activated at different time intervals, which may be a key to why the working memory of older adults is poorer. On the one hand the frontal area is thought to play a role in the active manipulation of information, acting as an attentional filter to determine which information will be selected for maintenance in working memory; while on the other hand the parietal cortex is involved in the actual maintenance of that information [48]. The reason why older adults activate the frontal areas less than younger ones, may be because this region reacts most sensitively to aging in terms of both structural and functional changes [49]; and therefore, another region takes over its role, which has a better preserved structural integrity.

According to the PBWM model, a gating mechanism enables the updating of working memory, and also ensures the protection of its content. *Gate opening* makes possible the entry of information into working memory and allows for its updating to occur. We found three components in the younger group, one of which was present in the older group as well, that significantly emerged from the EEG. When analysing *gate opening*, it is important to consider that this reflects the difference between reference switch and reference no-switch trials. Consequently, two important parameters do not equalize. One being that in the trial preceding the switch trials, the frame colour differs from the colour of the current frame, while in no- switch trials, the colour of the previous and currently viewed frame is the same. The other being their frequency difference: the probability of switch is 0.25, while the probability of no-switch trials is 0.75. Both are parameters that may influence the emergence of the ERP components.

The first component we observed is a negativity over the posterior regions between 150- 250 ms. In this range, various attentional effects are present, suggesting the possibility of selection negativity which distinctly influenced by the differential occurrence probabilities of the trial types [35,50,51]. Hence, it is possible that the emergence of the component was due to the differences between the two trial types and does not inherently characterise the *gate opening* process. This component had a larger amplitude in the older than the younger group, which suggests they were more involved in the selective analysis of the visual input.

The second component emerged between 300-500 ms and reached its positive maximum over the posterior areas. Given the similarity with the P3b component in oddball studies, differing probabilities may also influence this component [52,53]. This component may significantly contribute to the context updating-related P3b observed when the separation of subprocesses is omitted. Younger adults had a larger amplitude over the centro-parietal regions than the older ones, indicating emphasized processes in the former group.

The third component between 500-700 ms was also a positivity and had a parieto-occipital maximum. In the literature, the P600 component is known to occur over the posterior areas within this time window, encompassing both linguistic processes and executive functions [54]. It has also been found that the basal ganglia, which play a crucial role in gate opening in the PBWM model, modulate the P600 [55]. Alternatively, P600 belongs to the P3 family and is connected to cortical reorientation [56]. No age-related differences were found in the amplitude; however, larger activation was observed in extensive left-lateralized areas, mainly the orbitofrontal cortex and the lateral and medial temporal lobes in the younger group compared to the older one, in areas which have a role in: mapping contingencies [57]; object recognition [58]; consolidation and retrieval [59,60], respectively.

In summary, in *gate opening* not all conditions are balanced, so we cannot be certain that after the subtractions, we only see the effect of the subprocess emerging in the EEG. Nevertheless, we found age-related differences in all three evoked components, suggesting that this subprocess may operate with varying efficiency in the younger and older groups.

While it is important to be able to update the contents of working memory with new information, it is equally essential for efficient functioning to be able to protect it by ensuring that it remains intact against the influx of new information from external or internal sources. The protection of existing contents is made possible by *gate closing*. We found two components, similar in both groups, that characterized this process: a negativity above the occipital electrodes between 150-300 ms; and a frontal positivity between 300-600 ms. Considering that, as with *gate opening*, here also exist differences which are not eliminated by subtraction – in the comparison switch trial, the frame colour differs from the colour of the frame seen in the previous trial, while in the comparison no-switch trial, it matches; and the former occurs with a probability of 0.25, while the latter with 0.75 – so we can assume similar processes to those previously described for *gate opening*. However, the two age-groups showed no difference in the amplitude, scalp distribution or the source of this component.

The positivity between 300-600 ms had a frontal maximum in both age-groups; however, the cluster-based permutation analysis showed a significant cluster only in the older group. This is interesting for two reasons. Firstly, the involvement of frontal areas suggests that *gate closing* may be an active process, consistent with Nir-Cohen et al.’s [7] conception. Secondly, a common issue among older adults is their decreased ability to protect the contents of their working memory; so, they are very sensitive to the interference of task-irrelevant stimuli [e.g., 61,62]. We might assume that this phenomenon is connected to the *gate closing* process as irrelevant information can intrude if the *gate closing* is not effective enough. By contrast, we did not find any differences between younger and older adults in either the amplitude or the source of this component.

Summarizing the *gate closing* results, this subprocess was characterized by a posterior negativity between 150-300 ms and a frontal positivity between 300-600 ms, neither of which was sensitive to aging.

A limitation of the study is that the sLORETA method is not suitable for identifying activity in the basal ganglia, so not all sources that are part of the PBWM model can be identified. This is the reason why we remained at a descriptive level for source localization, as our EEG data can only provide indicative information in this regard.

## Summary

In this study our goal was to describe, for the first time, which ERP components are involved and how in the working memory subprocesses that can be disentangled using the reference back paradigm, as well as how aging affects these subprocesses. *Updating* was characterized by occipital negative components between 80-180 ms, 300-400 ms and 400-1,000 ms, with only the latter showing age-related differences. When analysing *substitution,* we observed a frontal positivity between 250-600 ms in younger adults, and a posterior positivity between 550-750 ms in older adults. In *gate opening* three parieto-occipital components emerged: a negativity between 150-250 ms, a positivity between 300-500 ms, and a positivity between 500-700 ms, all of which showed age-related differences. Finally, *gate closing* was characterised by an occipital negativity between 150-300 ms and a frontal positivity between 300-600 ms, neither of which changed between the two age groups.

Based on the above, we can conclude that the process of protecting information, i.e., *gate closing*, does not change with aging, contrary to what we might assume from older adults’ sensitivity to interference and inability to properly inhibit irrelevant information. Instead, *gate opening* is the process that is sensitive to age-related effects, and this is likely to be what leads to the different brain activity observed between the two age groups during *substitution*, i.e., the updating of working memory with new information.

## Acknowledgements

We thank Zsuzsanna D’Albini for her technical assistance and Nick Winnington-Ingram for language editing. This research was supported by the Hungarian Research Fund (OTKA K 115457 and 132880).

